# SCOUT: Single-cell outlier analysis in cancer

**DOI:** 10.1101/2020.03.25.007518

**Authors:** Giovana Ravizzoni Onzi, Juliano Luiz Faccioni, Alvaro G. Alvarado, Paula Andreghetto Bracco, Harley I. Kornblum, Guido Lenz

**Author notes:** These authors contributed equally to this work (Giovana Ravizzoni Onzi, Juliano Luiz Faccioni). Correspondence: *HIK; GL*.

## Abstract

Outliers are often ignored or even removed from data analysis. In cancer, however, single outlier cells can be of major importance, since they have uncommon characteristics that may confer capacity to invade, metastasize, or resist to therapy. Here we present the Single-Cell OUTlier analysis (SCOUT), a resource for single-cell data analysis focusing on outlier cells, and the SCOUT Selector (SCOUTS), an application to systematically apply SCOUT on a dataset over a wide range of biological markers. Using publicly available datasets of cancer samples obtained from mass cytometry and single-cell RNA-seq platforms, outlier cells for the expression of proteins or RNAs were identified and compared to their non-outlier counterparts among different samples. Our results show that analyzing single-cell data using SCOUT can uncover key information not easily observed in the analysis of the whole population.

## Introduction

Individual cells in a tumor are characterized by complex and dynamic profiles in gene expression and phenotypes that ultimately lead to treatment failure and disease recurrence. The combination of high parameterization and high throughput in single-cell techniques is essential for the detection and characterization of rare cells, as is the case of cancer stem cells or specific subpopulations of immune cells, that are present within the tumor mass and can be relevant for resistance to therapy (1, 2).

Outliers are observation points in a dataset that fall far outside the typically expected variation, occurring due to large discrepancy among observations in a group or due to experimental errors, which often leads to their a priori exclusion from data analysis (3). Nevertheless, cancer cells are especially varied and excluding outliers may impede the study of small subgroups or even individual cells with exceptional and important characteristics. To select and analyze these outliers we developed the Single-Cell OUTlier analysis (**SCOUT**, fig. 1A, see also **Sup. Mat. Video 1**). While SCOUT does not account for mutlivariate outlier analysis (4), it provides a more straightforward biological interpretation of the selected cells: outliers for a given marker can be directly linked to that marker’s metabolic implications.

**Fig. 1.**
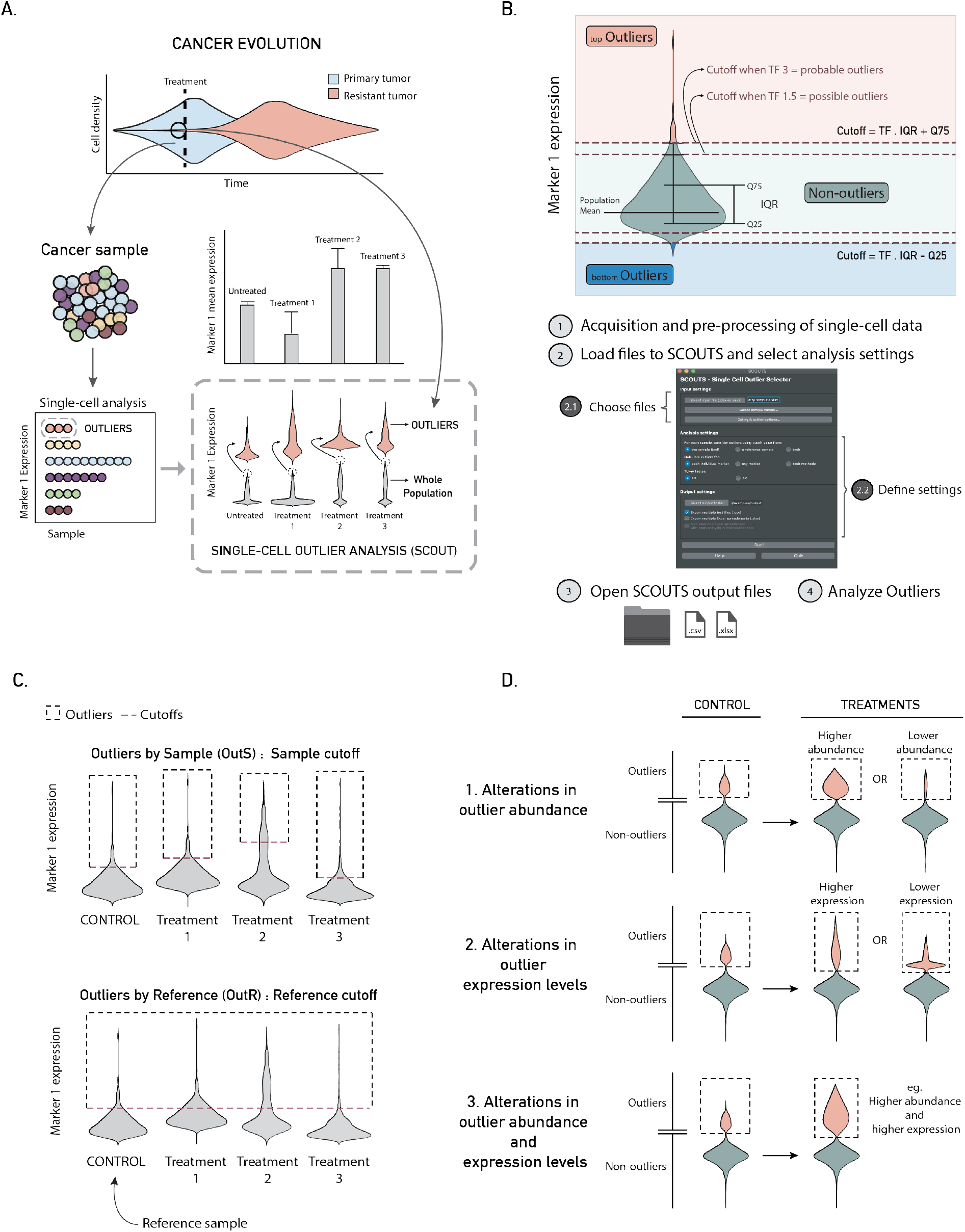
SCOUT and SCOUTS overview. **A**. Rare outlier cells with specific characteristics that are not reflected by the average population (bar plot) can represent key elements in cancer cell phenotypes or response to treatment. **B**. *Upper*: outlier detection by Tukey’s boxplot method. Dashed lines correspond to cutoff values either for top (orange) or bottom (blue) outliers. *Lower*: SCOUTS interface workflow. **C**. Outliers can be chosen by individual samples (OutS) or by a reference sample (OutR). **D**. Theoretical examples of possible alterations revealed by SCOUT when analyzing outliers. **TF**: Tukey factor; **IQR**: interquartile range; **Q25** first quartile; **Q75**: third quartile.

## How it works

To detect outlier cells, SCOUT uses the labeling method described by Tukey (5), also known as the boxplot method. Tukey’s method makes no distributional assumptions and does not depend on mean or standard deviation measures. Outliers are detected according to cutoffs established by:

i. determining the first and third quartiles of the sample;
ii. calculating the interquartile range (IQR);
iii. multiplying the IQR by a Tukey Factor (TF);
iv. summing the value obtained in the previous step to the third quartile.

The result is the cutoff for outliers at the top of the population distribution. Alternatively, one may subtract the value obtained in iii) from the first quartile, in order to obtain the cutoff for bottom outliers. The stringency of the detection depends on the value chosen for the TF; a TF of 1.5 detects *possible outliers,* while a TF of 3 detects *probable outliers* (fig. 1B, upper) (6). The resulting cells are deemed as *outliers for the given marker.*

In order to systematically select outlier cells from a population across multiple markers, we developed the Single-Cell OUTlier selector (**SCOUTS**). The outlier selection is performed according to settings defined by the user through a graphical user interface (GUI) (fig. 1B, lower). SCOUTS is a free, open-source software written in Python and distributed under the MIT license. For instructions on how to download and use SCOUTS, please read the Software availability section. See also **Sup Mat. Video 2**.

In SCOUTS, outliers can be defined as Outliers by Sample (**OutS**) - where outliers for each parameter are determined in each sample independently (fig. 1C, upper), or as Outliers by Reference (**OutR**), where a reference sample, such as a control sample, is used to determine the outlier cutoff for all other samples (fig. 1C, lower). Finally, possible results obtained when analyzing outliers identified by SCOUTS, such as alterations in their abundance or expression levels for a specific parameter, are illustrated in fig. 1D.

## Applications

We applied SCOUT to a publicly available mass cytometry dataset consisting of patient-derived melanoma samples treated with BRAF^V600E^ and MEK inhibitors (MEL-MC), containing paired samples of each patient before (Pre-Tx) and after four weeks of treatment (Week4) (7). In fig. 2A, heatmaps show the expression of different markers in the whole population, non-outliers and OutS for one of the patient samples. The *Out/Non-out* ratio, which corresponds to the ratio between the mean expression in outliers and nonoutliers, and *Relative* ratios, corresponding to ratios between treated and untreated samples, are also shown.

**Fig. 2.**
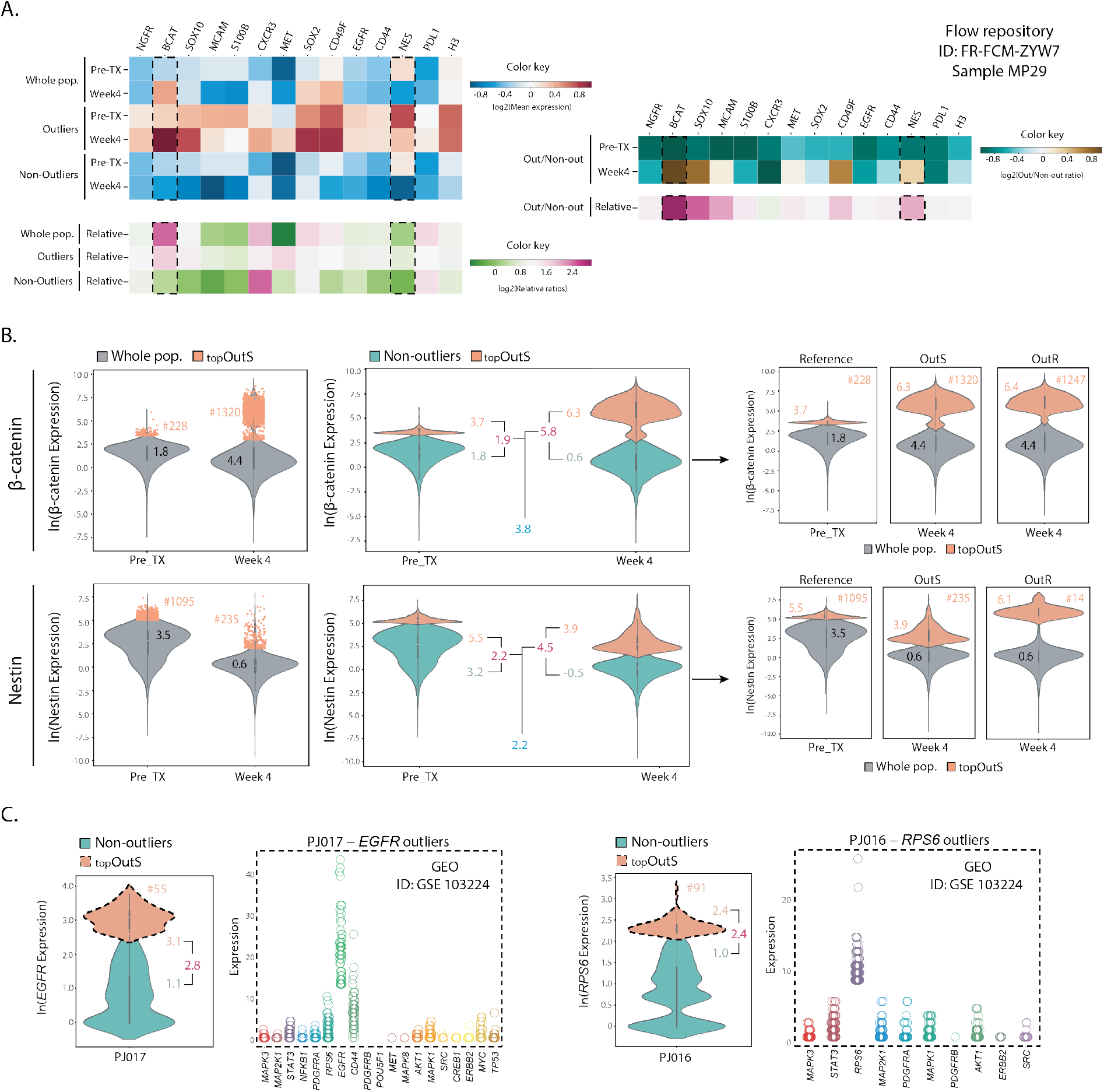
SCOUT applications using different single-cell datasets. **A**. Heatmaps of marker expression before (Pre-Tx) and after (Week4) treatment for sample MP29 of the MEL-MC dataset. Dashed black boxes denote specific markers selected for analysis. **B**. Violin plots of data shown in A. for β-catenin *(upper)* and Nestin *(lower)* outliers (OutS). Plots on the right present a comparison between OutS and OutR for each of these markers after treatment. **C**. SCOUT results for single-cell RNA-seq data. *Left*: violin and dot plots of *EGFR* expression and expression of other genes in *EGFR* outliers of patient PJ017. *Right*: violin and dot plots of *RPS6* expression and expression of other markers in *RPS6* outliers of patient PJ016. In Figures 2B and C, # indicates the absolute number of outliers. Black, orange and green numbers represent mean expression in the whole population, in the outlier population, and the non-outlier population, respectively. Numbers in red correspond to Out/Non-out ratios, while blue numbers represent Relative Out/Non-out ratios.

The comparisons between the *Relative* ratio of the whole population and the *Relative Out/Non-out* ratio demonstrate that SCOUT can uncover effects not observed in an average-based analysis of the total cell population. Taking the marker β-catenin as an example, there was an approximate 2.6-fold increase in the mean expression of the whole population after treatment, but this increase was due to a large increment in the number and expression levels of outliers only (fig. 2B, upper left and middle). The *Out/Non-out* ratios corroborated this finding, changing from 1.9 to 5.8 (fig. 2B, red), a 3.8- magnitude increase in natural logarithmic scale (approximate 50x linear increase). A difference between outliers and nonoutliers was also observed for markers like Nestin (**fig. 2B**, lower left), in which the mean expression in the whole population significantly decreased, yet the Out/Non-out ratio increased with treatment. Notably, both β-catenin and Nestin are known to be involved in pro-tumorigenic features, including drug resistance (8–10). OutS and OutR of β-catenin were very similar. Nonetheless, the differences between OutS and OutR of Nestin highlight the fact that there was a decrease in expression in the whole population with treatment and a consequent lower cutoff value in the Week4 sample, leading to a small amount of OutR when compared to OutS (fig. 2B, lower right).

SCOUT was next applied to single-cell RNA-seq data using a dataset of glioblastoma (GBM) patient-derived samples downloaded from GEO (11). Even though dealing with much smaller numbers of cells per sample (around 1,000 cells, 40x less than the mass cytometry dataset), SCOUT was able to detect outliers and reveal distinct patterns of gene expression in OutS of different samples. The PJ017 sample, for example, which had the *EGFR* gene amplified, had a high number of *EGFR* outliers. Notably, when we analyzed the expression of all other genes within the *EGFR* outliers we found that some genes had enriched expression in this subpopulation, such as *CD44* and *R0S6* (fig. 2C, left). The PJ016 sample, on the other hand, had a high number of *RPS6* outliers, with concomitant high expression levels of *STAT3* (fig. 2C, right). The example provided above indicates that there is potential value in examining multiple parameters within specific subsets of outliers that are established using only a single parameter. In this sense, it is noteworthy that integrative approaches of analysis can be performed in the outliers selected by SCOUT, such as the evaluation of expression correlations between markers in outliers and its comparison to correlations found in the whole population. These approaches may aggregate information on different targets that can be used to comprehend the biological implications of the outlier cells.

## Conclusion

Using SCOUT we were able to detect outliers for the expression of important molecules related to cancer progression and provide information on these rare cells that would be overlooked with other analytical approaches. To facilitate the application of this approach to data from different single-cell analysis platforms a GUI application, SCOUTS, was developed. Although caution should always be taken to avoid dealing with possible experimental artifacts, the study of outlier cells in cancer may contribute to the discovery of new therapeutic targets and aid in the design of more effective drug combinations.

## Material and methods

### SCOUTS: a GUI for applying SCOUT systematically

SCOUTS uses SCOUT to systematically select outliers as previously described. It accepts tabular data as input (either as a .csv or a .xlsx file), where the columns represent each marker, and rows represent the individual cells composing the population. The first row (header) must contain the names of the parameters, while the first column is reserved for cell and sample identification. We provide an example dataset on the project’s GitHub page.

Since sample naming conventions may vary widely between datasets, SCOUTS cannot automatically identify samples from a dataset only by reading the sample name column. To overcome this limitation, sample names must be individually input. SCOUTS allocates rows that contains the same user-defined text string as belonging to the same sample.

SCOUTS is able to select cells that are outliers for each marker, or all cells that are outliers for at least one marker. Additionally, it is also possible to select outliers at the bottom of the population distribution (low-expression outliers), or select that are not outliers. Furthermore, the user can choose to use one or both selection methods described in the main article: Outliers by Sample (OutS) and Outliers by Reference (OutR) (fig. 1C). The results are saved to a user-designated output folder in multiple files.

Ideally, the dataset should be preprocessed prior to SCOUTS. Since data preprocessing will depend on the nature of the experiment and analytical goals, we made the design choice of not to include preprocessing directly into SCOUTS’s workflow. However, SCOUTS provides the option of gating out poorly-stained cells (for mass cytometry data) and zero-read values (for scRNA-seq data).

### Test datasets

The MEL-MC mass cytometry dataset was downloaded from the FlowRepository database (ID: FR-FCM-ZYW7). The dataset consisted of melanoma samples from patients enrolled in a clinical trial in which biopsies were collected before and 4 weeks after treatment with targeted BRAF and MEK inhibitors (dabrafenib and trametinib, respectively) (7). Samples of three patients were chosen for analysis: MP29, MP34 and MP59 and, within these samples, only tumor cells that presented low (below the population median) *CD45* expression were considered. The GBM scRNA-seq dataset was downloaded from GEO (GEO accession GSE103224) and consisted of GBM patient-derived samples obtained from different donors (11).

The MEL-MC dataset was already normalized and gated (7), and we further selected the option provided in SCOUTS to exclude poorly stained cells, where for each cell the mean expression of all markers is calculated and cells with a mean value lower than 0.1 are considered as poorly stained and excluded from the analysis.

scRNA-seq results, due to the small amounts of starting material and low sampling, have inherently more noise than those obtained from bulk RNA-seq experiments (12) and, as a consequence, expression matrices usually contain excessive zero values. Therefore, for the GBM single-cell RNA-seq dataset, as a large number of zeros was observed for all samples, zero values were excluded to determine the cutoffs in the samples using the option provided in SCOUTS.

### Software availability

An overview of SCOUTS, as well as installation instructions, can be found at: www.ufrgs.br/labsinal/scouts.

The source code for SCOUTS is deposited at: https://github.com/jfaccioni/scouts.

For a more in-depth explanation of how SCOUTS works, please read the full documentation at: https://scouts.readthedocs.io/en/master/.

## Supporting information

Sup. Mat. Video 1

Sup. Mat. Video 2

## ACKNOWLEDGEMENTS

We acknowledge the support provided by the Flow Cytometry Core of UCLA and thank Alejandro Garcia for the help with mass cytometry experiments. We also thank Rodrigo Allende for the video recordings. This work was supported by Conselho Nacional de Desenvolvimento Científico e Tecnológico CNPq, Universal (475882/ 2012-1) and Novas Terapias Portadoras de Futuro (457394/2013-7);PROBITEC CAPES (907/2012); ICGEB (405231/2015-6); and The Dr. Miriam and Sheldon G. Adelson Medical Research Foundation and the UCLA SPORE in Brain Cancer (P50 CA211015) (HIK). G.R.O was recipient of a doctorate fellowship from CNPq (Conselho Nacional de Desenvolvimento Científico e Tecnológico) and of a sandwich doctorate fellowship from CAPES (Coordenação de Aperfeiçoamento de Pessoal de Nível Superior). J.F. is recipient of a doctorate fellowship from CNPq. P.A.B was recipient of doctorate fellowship from CAPES. G.L. is recipient of CNPq research productivity fellowship.

